# R-spondin2 regulates planar cell polarity in the vertebrate neural plate

**DOI:** 10.64898/2026.05.22.727270

**Authors:** Ilya Chuykin, Sergei Y. Sokol

## Abstract

Vertebrate neural tube closure requires planar cell polarity (PCP) signaling to coordinate cell behaviors in the neuroepithelium. In the *Xenopus* neural plate, PCP is marked by enrichment of the core PCP protein Vangl2 at the anterior edge of every cell, but the distribution of the extracellular factors modulating this asymmetry remains largely unknown. Here, we identify the secreted protein R-spondin2 (Rspo2) as a modulator of neural plate PCP. Rspo2 exhibits predominantly anterior localization in neuroepithelial cells. Morpholino-mediated knockdown of Rspo2 causes neural tube closure defects and disrupts the anterior enrichment of Vangl2. Rspo2 associates with Vangl2 and inhibits FGF receptor-dependent Vangl2 tyrosine phosphorylation *in vivo*, as shown by both depletion and overexpression experiments. Rspo2 domain analysis shows that the thrombospondin domain contributes to both anterior Rspo2 membrane enrichment and inhibition of Vangl2 phosphorylation. Together, these findings identify a role of Rspo2 in PCP signaling in the *Xenopus* neural plate and support a model in which anteriorly localized Rspo2 helps maintain Vangl2 asymmetry while limiting Vangl2 tyrosine phosphorylation during neural plate morphogenesis.

## INTRODUCTION

Neural development depends on signaling networks that coordinate cell fate, spatial patterning and morphogenetic behaviors that shape the central and peripheral nervous systems in vertebrate embryos. Among these, the planar cell polarity (PCP) pathway orients neuroepithelial cells along the anteroposterior axis and is essential for neural tube closure (Butler and Wallingford, 2017; Nikolopoulou et al., 2017). At the cellular level, PCP is established through the asymmetric organization of core protein complexes: Van Gogh-like and Prickle (Vangl-Pk) and Frizzled and Dishevelled (Fz-Dvl), with Flamingo/Celsr bridging both (Adler, 2012; Basta et al., 2025; Devenport, 2014; Goodrich and Strutt, 2011; Weiner et al., 2025). The developmental importance of PCP is highlighted by genetic studies linking mutations in pathway components to neural tube closure defects (Ciruna et al., 2006; Jessen et al., 2002; Kibar et al., 2001; Lu et al., 2004; Murdoch et al., 2001; Wang et al., 2006; Ybot-Gonzalez et al., 2007).

The localization of core PCP complexes is believed to be established by the Wnt and FGF pathways, possibly via the phosphorylation state of the core PCP protein Vangl2 (Chu and Sokol, 2016; Chuykin et al., 2021; Chuykin and Sokol, 2025; Gao et al., 2018; Gao et al., 2011; Kelly et al., 2016; Ossipova et al., 2015; Strutt et al., 2019; Wu et al., 2013; Yang et al., 2017). However, how these signals are distributed within the neural plate to coordinate PCP is less clear. Notably, Wnt11 was recently reported to exhibit polarized distribution in individual Xenopus neuroepithelial cells (Mii et al., 2026). These findings raise the possibility that other secreted factors may also localize asymmetrically to compose a new extracellular layer of PCP regulation.

R-spondins (Rspo1-4) are conserved secreted proteins that regulate embryonic development and tissue homeostasis. R-spondins share common domain organization, including two N-terminal furin-like (Fu) cysteine-rich domains, a thrombospondin type 1 (TSP1) domain, and a basic C-terminal region (de Lau et al., 2014; Jin and Yoon, 2012). Rspo proteins are known to promote Wnt/β-catenin signaling by acting through LGR4/5/6 receptors to inhibit the E3 ubiquitin ligases RNF43/ZNRF3 and thereby stabilize Frizzled receptors at the cell membrane (Carmon et al., 2011; de Lau et al., 2011; Hao et al., 2012; Koo et al., 2012). However, subsequent studies revealed that Rspo proteins, especially Rspo2 and Rspo3, can potentiate Wnt/β-catenin signaling in the absence of LGR4/5/6 (Lebensohn and Rohatgi, 2018; Park et al., 2018; Szenker-Ravi et al., 2018) and interact with other surface molecules to regulate embryonic development (Dubey et al., 2020; Niehrs et al., 2024).

Rspo2 has essential developmental roles in the limb, lung, and craniofacial development (Aoki et al., 2008; Bell et al., 2008; Jin et al., 2011; Nam et al., 2007; Yamada et al., 2009). Notably, these processes also depend on PCP and FGF signaling (Gao et al., 2018; Gao et al., 2011; Min et al., 1998; Peters et al., 1994; Sekine et al., 1999; Yates et al., 2010). These observations suggest that Rspo2 may crosstalk with the FGF- and PCP-dependent morphogenetic programs. Although Rspo2 was initially identified as a Wnt/β-catenin activator (Kazanskaya et al., 2004), later work showed that it modulates both Wnt and FGF signaling in a context-specific manner (Reis and Sokol, 2020; Reis and Sokol, 2021). A recent study reported that Rspo2 is the only member of the Rspo family that interacts with FGFR4 (Lee et al., 2024). Of note, a closely related R-spondin, Rspo3, interacts with Syndecan4 and is required for body axis elongation in *Xenopus* embryos, suggesting a link to PCP signaling (Ohkawara et al., 2011). These observations, together with our recent finding that FGF signaling intersects with PCP during neural tube closure (Chuykin and Sokol, 2025), raise the possibility that Rspo2 might be involved in PCP signaling.

To test this hypothesis, we investigated Rspo2 localization and function in the vertebrate neuroectoderm. The *Xenopus* neural plate offers a tractable vertebrate model for PCP, because Vangl2 localizes asymmetrically to anterior cell boundaries, and this distribution depends on its interaction with Prickle proteins (Jenny et al., 2003; Ossipova et al., 2015). Our experiments show planar polarization of Rspo2 at the anterior edges of neuroepithelial cells, its requirement for Vangl2 accumulation, and the role of Rspo in FGF-dependent Vangl2 phosphorylation.

## RESULTS AND DISCUSSION

### Planar polarization of Rspo2 in the Xenopus neuroepithelium

Rspo2 is abundantly expressed in *Xenopus* neurulae, with prominent expression in the anterior neural plate and lateral plate mesoderm (Kazanskaya et al., 2004), raising the possibility that it contributes to PCP-dependent neuroepithelial morphogenesis. We therefore evaluated Rspo2 localization in neural plate cells. To address this, we injected low doses of RNA encoding tagged Rspo2 into a dorsal animal blastomere to target the neural plate and examined protein distribution in neurula-stage embryos by immunostaining. Rspo2 was detected in discrete membrane-associated puncta that were preferentially enriched at anterior cell boundaries, rather than being uniformly distributed along cell boundaries **(Fig. 1A-B′, arrows)**. Quantitative analysis of polarity vectors confirmed a strong anterior bias in Rspo2 localization **(Fig. 1C)**. Rspo2-GFP fused to a different tag exhibited similar localization in the neural plate (data not shown). These observations indicate that Rspo2 can polarize in the plane of the neuroepithelium.

**Figure 1.**
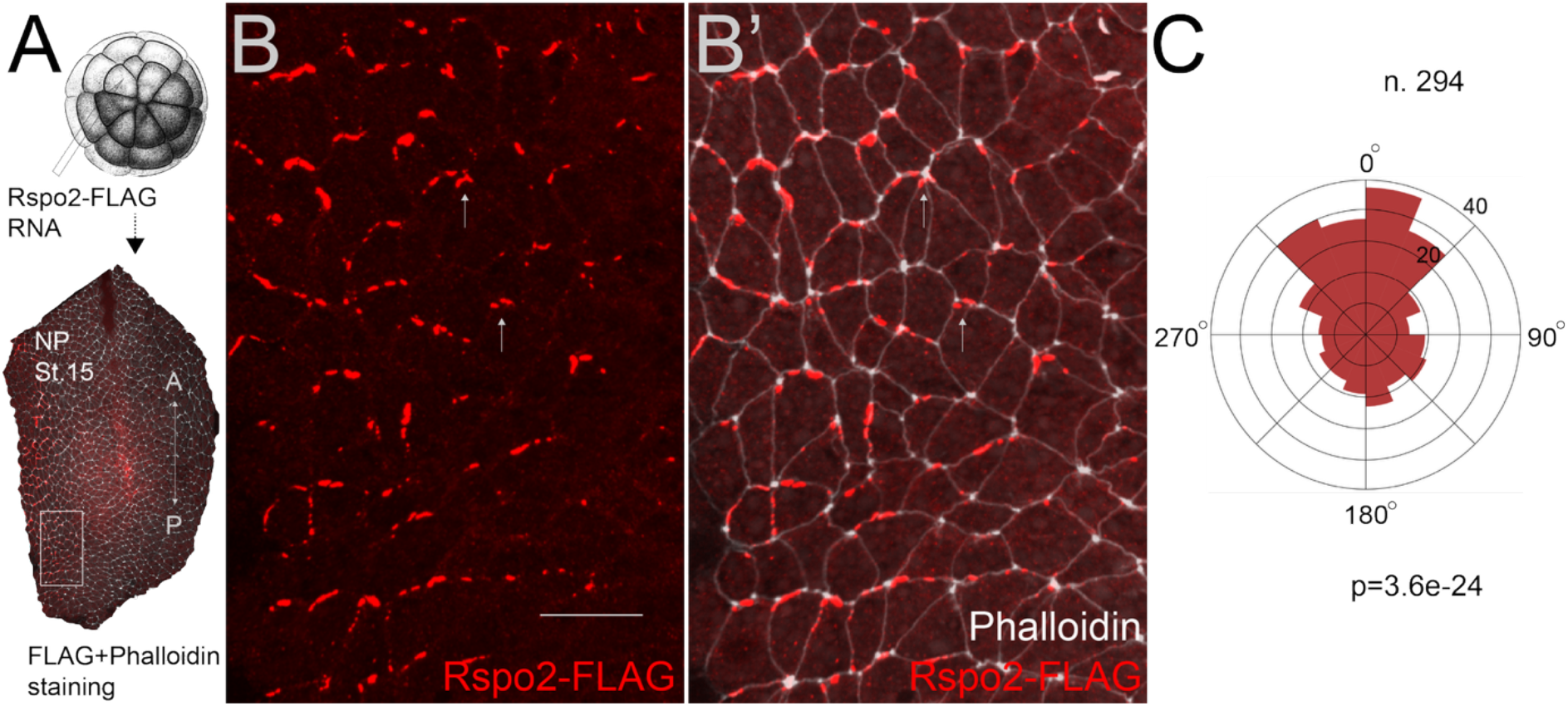
R-spondin2 displays anteriorly biased membrane localization in the neuroepithelium. (A) Experimental scheme. *Xenopus* embryos were injected with RNA encoding Rspo2-FLAG, 80 pg, into one dorsal animal blastomere at the 32-cell stage, developed until neurula stage 15 (st.15) and subjected to FLAG immunostaining. Neural plates (NP) were imaged *en face*. Anteroposterior axis (AP) is indicated. The boxed region indicates the area analyzed at higher magnification in B and B′. (B, B′) Representative images of Rspo2-FLAG localization in the neuroectoderm. Rspo2-FLAG accumulates in discrete membrane-associated puncta and is preferentially enriched at anterior cell boundaries. Rspo2-FLAG is shown alone in red (B) and merged with phalloidin staining marking cell boundaries in light grey (B′). Arrows indicate examples of anteriorly enriched Rspo2-FLAG puncta. (C) Quantification of Rspo2-FLAG polarity shown as a rose plot of polarity angles. Each sector represents the number of cells with membrane-associated Rspo2-FLAG enrichment in a given direction, with 0° corresponding to anterior. The pooled distribution of cells from three embryos showed a strong anterior bias (n = 294 cells; Rayleigh test, p=3.6*10^-24^). Cell outlines were segmented from the phalloidin channel using Cellpose, and polarity vectors were calculated from membrane-associated Rspo2-FLAG intensity using custom Python scripts. Scale bar, 30 µm.

### Rspo2 depletion impairs neural tube closure

The anterior enrichment of Rspo2 suggests a potential role in PCP signaling in the neural plate (Nikolopoulou et al., 2017). We therefore tested whether targeted Rspo2 depletion affects neural tube closure in *Xenopus* embryos. Embryos unilaterally injected with Rspo2 MO exhibited a broad range of neural tube closure defects **(Fig. 2A-C)**. Quantification revealed a marked increase in the frequency of neural tube defects in Rspo2 MO-injected embryos compared with uninjected controls **(Fig. 2D)**.

**Figure 2.**
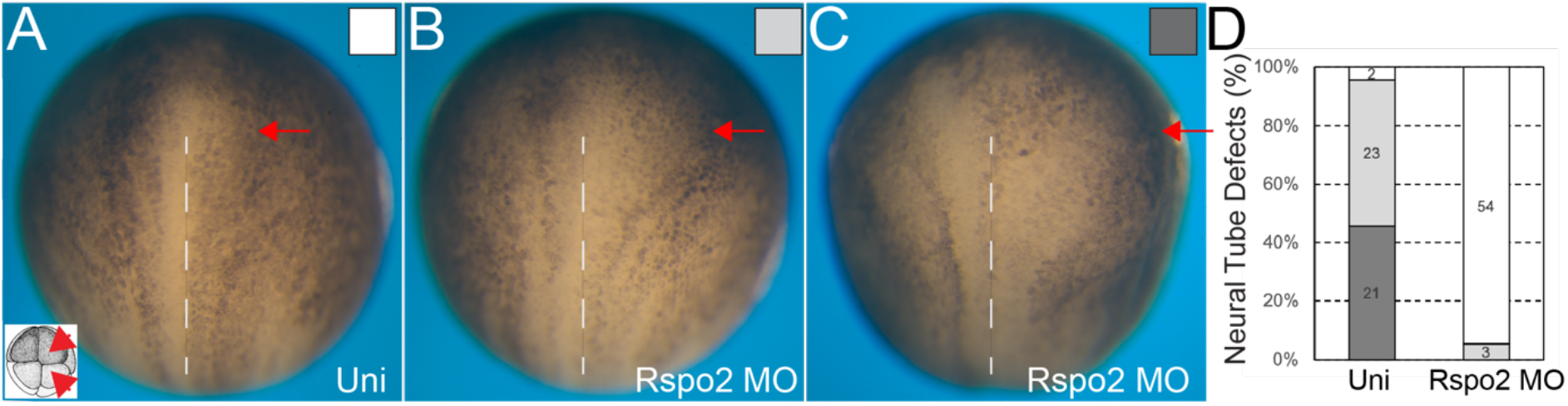
R-spondin2 depletion leads to neural tube defects (NTDs) in *Xenopus* embryos. *Xenopus* embryos were unilaterally injected with R-spondin2 morpholino (Rspo2 MO) into one dorsal and one ventral animal blastomere at the 8-cell stage (A, inset). Embryos were fixed at stages 16-17, and neural tube closure phenotypes were classified as none (A), mild (B), or severe (C), indicated by white, light grey, and dark grey squares, respectively. Representative embryos are shown (A-C). The position of neural fold is indicated by red arrow at the injected side, and the embryonic midline is indicated by a dotted line. (D) Quantification of neural tube defects. Frequencies of mild and severe NTDs are shown for uninjected (Uni) and Rspo2 MO embryos. Numbers of scored embryos per group are indicated within bars. Data are representative of two independent experiments.

To ensure that neural tissue specification was not affected, we performed immunostaining for Sox3, a pan-neural marker (Collignon et al., 1996; Penzel et al., 1997). Sox3 expression was comparable between injected and uninjected regions of the neural plate in embryos injected with either Rspo2 or control MOs **(Fig. 3A-B′)**. These results suggest that, under these conditions, the neural tube closure defects in Rspo2 morphants primarily reflect impaired morphogenesis rather than altered neural specification.

**Figure 3.**
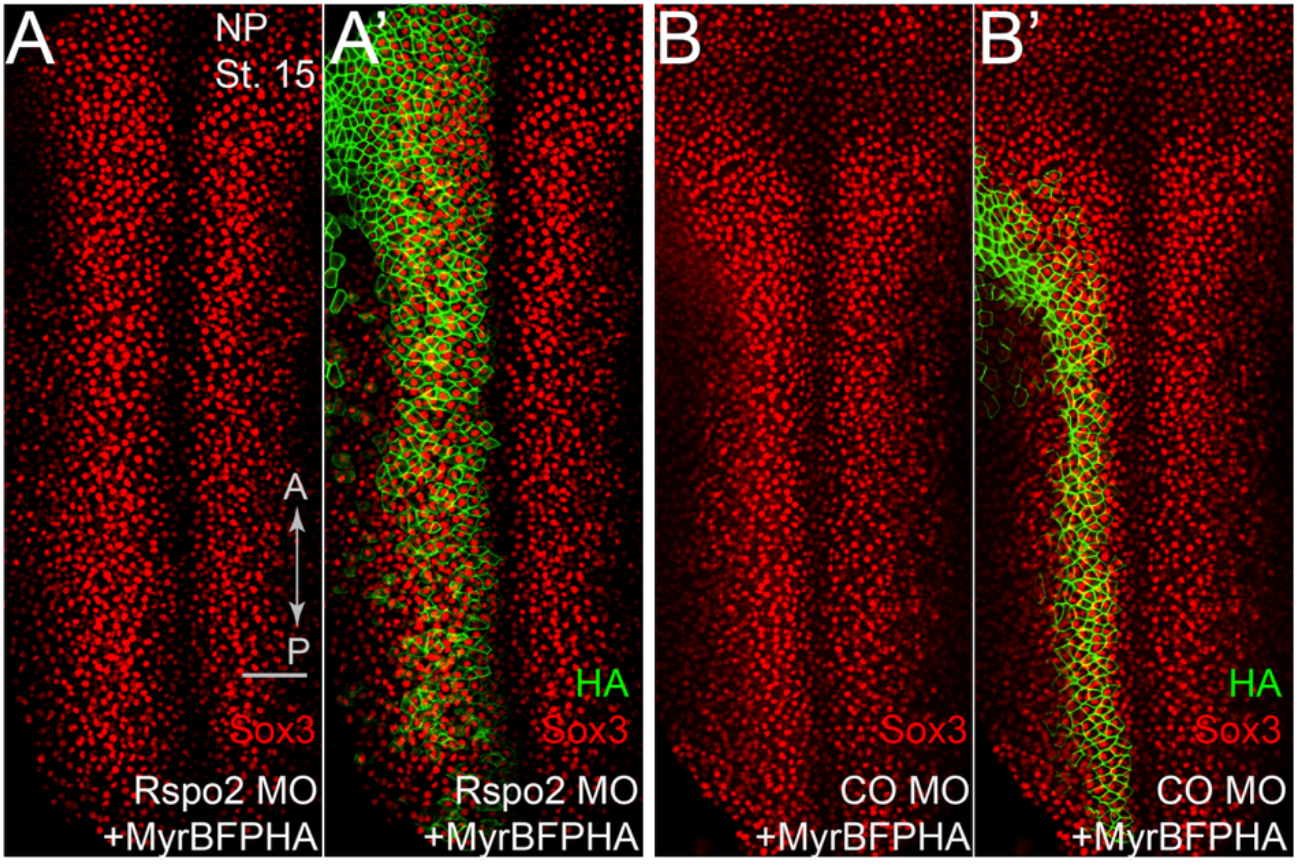
Sox3 expression is maintained in Rspo2 morphants. Eight-cell *Xenopus* embryos were co-injected with 10 ng of R-spondin2 splice-blocking morpholino (Rspo2 MO) or control morpholino (Co MO, 10 ng), together with Myr-BFP-HA RNA (40 pg) as a lineage tracer into one dorsal animal blastomere. Embryos were fixed at stage 15 (st.15) and co-immunostained for Sox3 (red) and HA (green). The anteroposterior (A-P) axis is indicated. Top-view neural plate images are shown. (A, A′) Rspo2 MO-injected embryos; (B, B′) control MO-injected embryos. Panels (A, B) show Sox3 staining, and (A′, B′) show merged images (HA+Sox3). Sox3 expression appears comparable between injected and uninjected regions in both Rspo2 morphants and controls. Scale bar, 200 µm.

Although neural tube defects have not been reported as a major phenotype in Rspo2 mutant mice (Bell et al., 2008), Rspo2 may still have context-dependent roles in neurulation that could be compensated in stable mutant backgrounds.

### Rspo2 is required for Vangl2 planar polarity in the neuroepithelium

To test whether Rspo2 contributes to PCP in the neural plate, we examined Vangl2 localization in Rspo2 morphant cells. Vangl2 is a core PCP protein that normally accumulates at anterior cell boundaries in the *Xenopus* neuroepithelium (Ossipova et al., 2015). In control regions of the neural plate, Vangl2 displayed clear asymmetry, with preferential enrichment at anterior cell junctions **(Fig. 4A, B, B′, arrows)**. By contrast, in Rspo2 MO-injected cells, Vangl2 localization was less polarized and more uniformly distributed along cell membranes **(Fig. 4B, B′, asterisks)**. Quantitative analysis of polarity vectors confirmed reduced alignment along the anteroposterior axis in Rspo2-depleted cells **(Fig. 4C, D)**. The disruption of Vangl2 asymmetry in Rspo2 morphants supports a role for Rspo2 in PCP regulation in the neural plate.

**Figure 4.**
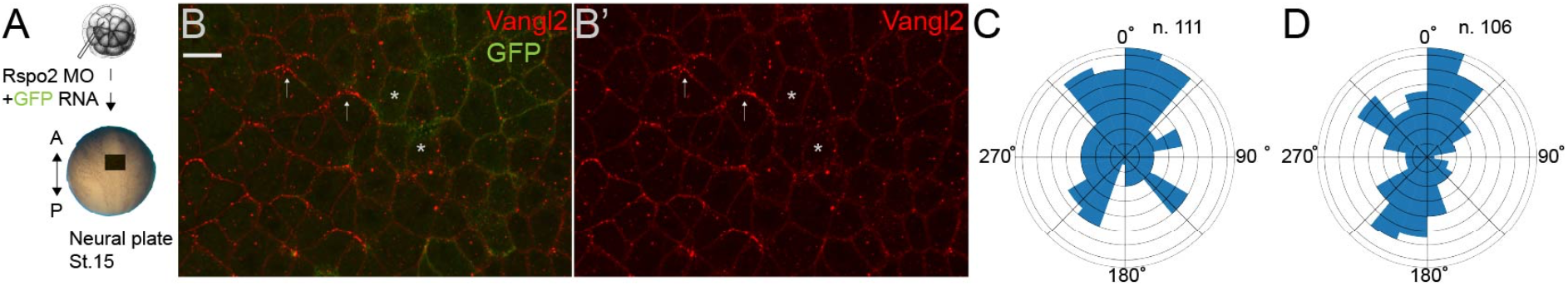
Vangl2 planar asymmetry is perturbed in Rspo2 morphants. (A) Experimental scheme. *Xenopus* embryos were injected at the 32-cell stage with 5 nl of R-spondin2 splice blocking morpholino (Rspo2 MO), 10 ng, into one dorsal animal blastomere together with GFP RNA as a lineage tracer. Embryos were collected and fixed at neurula stage 15 (st.15), and immunostained for Vangl2. The anteroposterior (AP) axis is indicated. (B, B′) Representative *en face* images of the neural plate showing GFP-positive cells (green) and Vangl2 localization (red). Vangl2 is enriched at anterior cell junctions in control regions (arrows). Asterisks mark GFP-positive morphant cells. (C, D) Quantification of Vangl2 planar polarity shown as rose plots of polarity angles for control (C) and Rspo2 MO-injected (D) neuroectoderm. 0° corresponding to anterior. Each sector represents the number of cells exhibiting membrane-associated Vangl2 enrichment in a given direction. Polarity was quantified by measuring the angle of a vector drawn from the cell centroid to the intensity-weighted centroid of Vangl2 signal restricted to the cell membrane. Numbers of analyzed cells (n) are indicated.

### Rspo2 associates with Vangl2 and attenuates FGFR-induced Vangl2 tyrosine phosphorylation

Since both Rspo2 and Vangl2 were localized to anterior cell sides, we next asked whether the two proteins are present in the same protein complex. Vangl2 was pulled down together with Rspo2 indicating the physical association between these proteins **(Fig. 5A)**. This observation suggests that Rspo2 may interact with Vangl2 directly or indirectly, potentially through a coreceptor, at anterior cell me mbranes.

**Figure 5.**
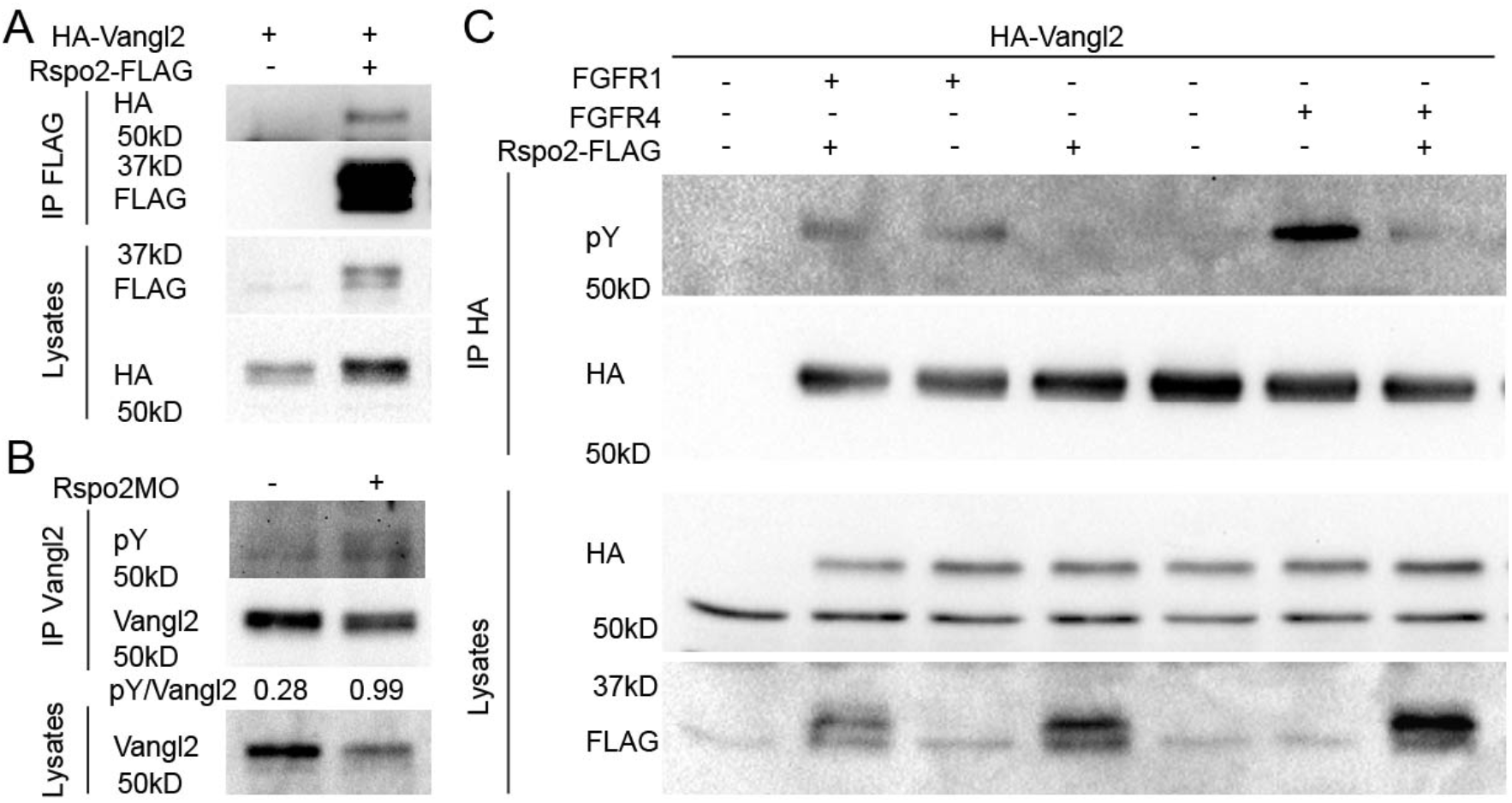
Rspo2 associates with Vangl2 and attenuates FGFR4-dependent Vangl2 tyrosine phosphorylation. **(A, B)** Animal cap explants were dissected at stages 8-9, cultured until sibling embryos reached the indicated stages, and processed for immunoprecipitation and immunoblotting. **(A)** Four-to eight-cell *Xenopus* embryos were injected with RNAs encoding HA-Vangl2, with or without Rspo2-FLAG, as indicated. Explants were cultured until sibling embryos reached stage 12 and subjected to immunoprecipitation with anti-FLAG agarose. Co-immunoprecipitated HA-Vangl2 was detected by immunoblotting with anti-HA antibody. **(B)** Explants from control embryos or embryos injected with translation-blocking Rspo2 morpholino, 10 ng, were cultured until stage 17 and subjected to immunoprecipitation with rabbit polyclonal anti-Vangl2 antibody followed by immunoblotting with anti-phosphotyrosine antibody. Increased Vangl2 phosphorylation was detected in Rspo2 morphant explants. **(C)** Four-to eight-cell embryos were injected with RNA encoding HA-Vangl2 together with FGFR1 or FGFR4 constructs, in the presence or absence of Rspo2-FLAG, as indicated. Embryos were lysed at stage 12 and subjected to immunoprecipitation with HA-Trap beads. Rspo2 attenuates FGFR4-, but not FGFR1-induced, Vangl2 tyrosine phosphorylation. Pulldowns and lysates were immunoblotted with anti-phosphotyrosine, anti-HA, anti-FLAG, and anti-Vangl2 antibodies, as indicated.

Because Rspo2 has been shown to antagonize FGF pathway activity (Reis and Sokol, 2020), and we found that FGFR signaling promotes Vangl2 tyrosine phosphorylation (Chuykin and Sokol, 2025), we assessed whether Rspo2 modulates FGFR-dependent Vangl2 phosphorylation. Rspo2 depletion increased endogenous Vangl2 tyrosine phosphorylation in ectodermal explants, suggesting that Rspo2 normally limits Vangl2 phosphorylation in embryonic tissues **(Fig. 5B)**.

Vangl2 tyrosine phosphorylation was previously shown to require FGFR1 (Chuykin and Sokol, 2025). However, both FGFR1 and FGFR4 are expressed in the neural plate (Lea et al., 2009; Yamagishi and Okamoto, 2010) and may, therefore, be involved in Vangl2 phosphorylation. To test the effect of Rspo2 on both receptors, we coexpressed Rspo2, Vangl2 and either FGFR1 or FGFR4. These gain-of-function studies showed that Rspo2 strongly attenuated FGFR4-induced but not FGFR1-induced Vangl2 tyrosine phosphorylation **(Fig. 5C)**, consistent with the reported preferential interaction between Rspo2 and FGFR4 (Lee et al., 2024).

These experiments suggest that Rspo2 negatively regulates FGFR4-dependent Vangl2 phosphorylation in embryos. This could involve heteromeric FGFR1-FGFR4 receptor complexes, as described for other receptor tyrosine kinases (Campana et al., 2025), although this possibility remains to be tested.

### The TSP1 domain contributes to Rspo2 anterior membrane enrichment and inhibition of Vangl2 phosphorylation

We next asked which domains of Rspo2 are required for its polarized distribution in the NP and activity. Full-length Rspo2-GFP showed anteriorly enriched membrane-associated puncta **(Fig. 6A, B, B′)**. Deletion of the furin-like domains did not disrupt this pattern, as Rspo2ΔF retained anterior membrane localization **(Fig. 6C, C′)**. In contrast, Rspo2ΔT, which lacks the TSP1 domain, did not show obvious anterior membrane enrichment and appeared more diffusely distributed **(Fig. 6D, D′)**. These observations suggest that the TSP1 domain, but not the furin-like domains, contributes to anterior membrane enrichment of Rspo2 in the neuroepithelium.

**Figure 6.**
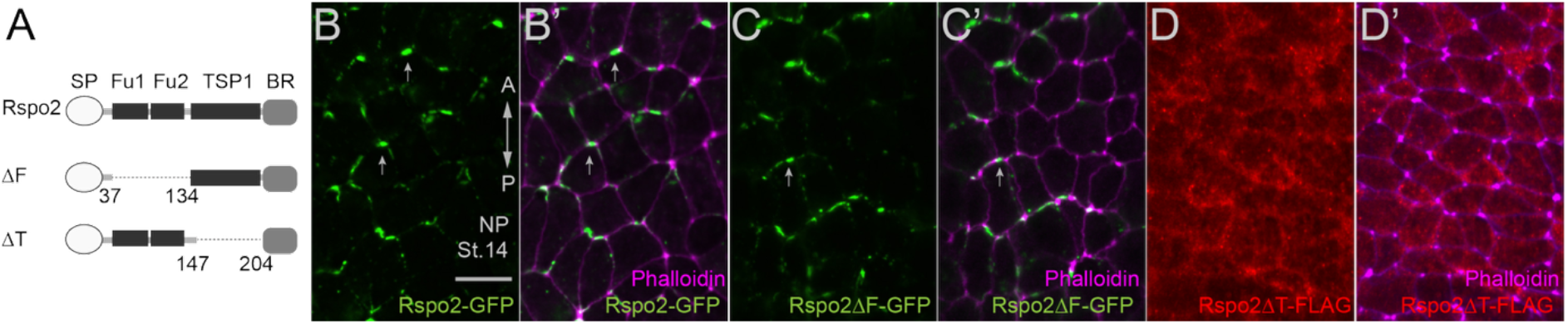
The thrombospondin domain contributes to anterior membrane enrichment of Rspo2 in the neuroepithelium. *Xenopus* embryos were injected at the 16-to 32-cell stage into one dorsal animal blastomere with RNAs encoding the indicated Rspo2 constructs and analyzed in the neural plate (NP) at neurula stage 14 (st. 14). Neural plates were imaged *en face* with the anteroposterior (AP) axis oriented vertically. **(A)** Schematic representation of full-length Rspo2 and deletion constructs used in this study. SP, signal peptide; Fu1 and Fu2, furin-like domains; TSP1, thrombospondin type 1 domain; BR, basic region. Amino acid positions delimiting the deleted regions are indicated. **(B-C′)** Representative images of full-length Rspo2-GFP and Rspo2ΔF-GFP, lacking the furin-like domains. Both constructs accumulate in membrane-associated puncta and show preferential enrichment at anterior cell boundaries. GFP fluorescence is shown alone in green **(B, C)** and merged with phalloidin staining in magenta **(B′, C′). (D, D′)** Representative images of immunostained Rspo2ΔT-FLAG, lacking the thrombospondin domain. In contrast to full-length Rspo2 and Rspo2ΔF, Rspo2ΔT-FLAG does not show obvious anterior membrane enrichment and appears more diffusely distributed. The FLAG signal is shown alone in red (**D**) and merged with phalloidin staining in magenta (**D′**). Scale bar, 30 µm.

We then tested whether the same domain was required for the inhibitory effect of Rspo2 on Vangl2 tyrosine phosphorylation. Full-length Rspo2 reduced FGFR4-induced Vangl2 tyrosine phosphorylation, and Rspo2ΔF retained this inhibitory activity **(Fig. 7A)**. In contrast, deletion of the TSP1 domain impaired the ability of Rspo2 to suppress Vangl2 phosphorylation **(Fig. 7A)**.

**Figure 7.**
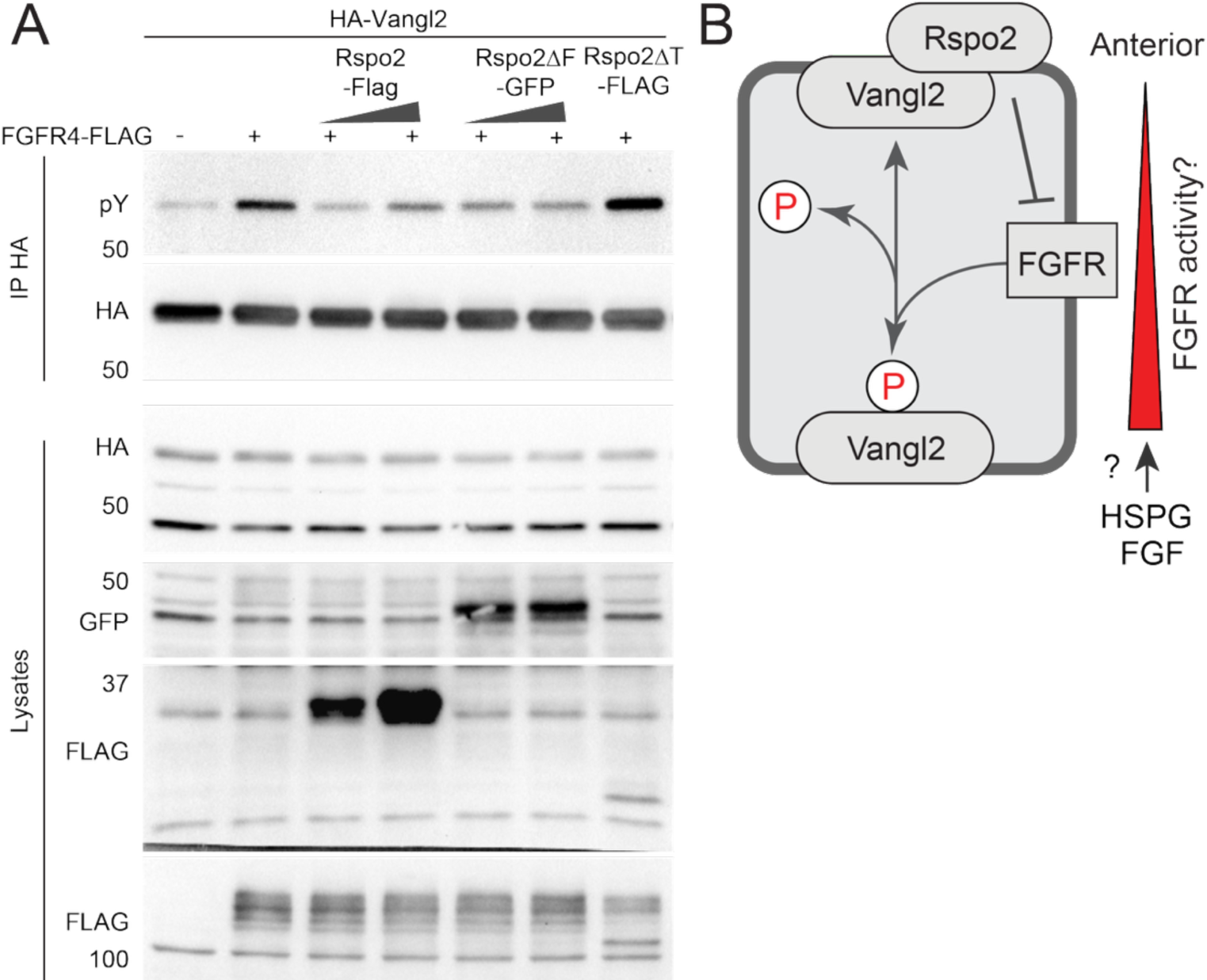
The thrombospondin domain of Rspo2 contributes to inhibition of Vangl2 tyrosine phosphorylation. (A) Four to eight cell *Xenopus* embryos were injected with RNAs encoding HA-Vangl2, 50 pg, FGFR4-FLAG, 100 pg with or without 250 or 500 pg of RNAs encoding full-length Rspo2-FLAG, Rspo2ΔF-GFP, or Rspo2ΔT-FLAG, as indicated. Embryos were lysed at stage 12 and subjected to immunoprecipitation with HA-trap beads, followed by immunoblotting with anti-phosphotyrosine (pY) antibody to assess Vangl2 tyrosine phosphorylation. Rspo2 and Rspo2ΔF, but not Rspo2ΔT, inhibit FGFR4-induced Vangl2 tyrosine phosphorylation. Pulldowns and lysates were immunoblotted with anti-pY, anti-HA, anti-GFP, or anti-FLAG antibodies, as indicated, to detect protein expression. (B) Model. Anteriorly localized Rspo2 limits FGFR-dependent Vangl2 tyrosine phosphorylation at anterior cell boundaries, resulting in higher FGFR activity at posterior boundaries. Spatial restriction of the Vangl2 phosphorylation by available FGF ligands or heparan sulfate proteoglycans may contribute to neural plate PCP.

Our findings highlight the TSP1 domain as a key region for further mechanistic analysis of Rspo2 function in PCP. Future studies should define smaller motifs within the TSP1 domain that mediate Rspo2 activity in PCP, identify the motif required for Vangl2 binding, and determine whether Rspo2, Vangl2, and FGFR4 can assemble into a common protein complex. It will also be important to address the potential contribution of the C-terminal basic region to Rspo2 function, because the TSP/basic region of R-spondins can mediate interactions with heparan sulfate proteoglycans (Dubey et al., 2020).

Together, our findings identify Rspo2 as a regulator of PCP in the *Xenopus* neural plate. Rspo2 is enriched anteriorly, is required for Vangl2 planar polarity, associates with Vangl2, and limits FGFR4-dependent Vangl2 tyrosine phosphorylation. These observations support a model in which anteriorly localized Rspo2 contributes to PCP by locally modulating FGFR-dependent effects on Vangl2, including its tyrosine phosphorylation **(Fig. 7B)**. This mechanism may influence the spatial regulation of Vangl2 complex assembly along the anteroposterior axis, although whether localized tyrosine phosphorylation directly controls Vangl2 planar asymmetry remains to be tested. Future studies defining the planar distribution and functional roles of FGF ligands, FGFRs, and heparan sulfate proteoglycans such as Syndecan4 (Escobedo et al., 2013; Ohkawara et al., 2011) will clarify how extracellular cues are localized to regulate PCP in the neural plate.

## METHODS

### Plasmids, morpholinos, and mRNA synthesis

Plasmids encoding HA-tagged *Xenopus laevis* Vangl2 (Chu and Sokol, 2016), mouse FGFR1-3xFLAG (Brewer et al., 2015), human FGFR4-V5-His (Gudernova et al., 2016), GFP (Ossipova et al., 2015), and myr-BFP-HA (Matsuda et al., 2023) were described previously. FGFR4-V5-FLAG construct was generated by subcloning. Rspo2 constructs used in this study included full-length Rspo2-FLAG, Rspo2-FLAG-GFP, Rspo2ΔF-GFP, and Rspo2ΔT-FLAG (Reis and Sokol, 2020). The Rspo2ΔF lacks the furin-like domain (aa 37-134), whereas Rspo2ΔT lacks the TSP domain (aa 147-204). Capped mRNAs were synthesized from linearized plasmids using the mMessage mMachine kit (Ambion) according to the manufacturer’s instructions. RNA amounts used for individual experiments are indicated in the figure legends. Rspo2 knockdown was performed using previously described translation-blocking or splice-blocking Rspo2 morpholino oligonucleotides (Reis and Sokol, 2020) with the following sequences: RspoMO^ATG^, 5′-AAAGAGTTGAAACTGCATTTGG-3′; RspoMO^SB^, 5′-GCAGCCTGGATACACAGAAACAAGA-3′. A standard control morpholino (Gene Tools) was used where indicated and had the sequence 5′-GCTTCAGCTAGTGACACATGCAT-3′.

### *Xenopus* embryo culture and microinjections

All animal procedures were performed in accordance with institutional guidelines and were approved by the Institutional Animal Care and Use Committee (IACUC) at the Icahn School of Medicine at Mount Sinai. *Xenopus laevis* eggs were fertilized *in vitro* and cultured in 0.1× Marc’s Modified Ringer’s solution (Chuykin et al., 2021). Embryos were staged according to Nieuwkoop and Faber (Nieuwkoop and Faber, 1967). For microinjections, embryos were transferred to 3% Ficoll in 0.6× MMR and injected with 5-10 nl of solution containing mRNA and/or morpholino oligonucleotides (MOs). For analysis of Vangl2 planar polarity, embryos were injected at the 32-cell stage into one dorsal animal blastomere with Rspo2 MO together with GFP RNA as a lineage tracer. For neural tube closure assays, embryos were injected unilaterally at the 8-cell stage into one dorsal and one ventral animal blastomere with Rspo2 MO. For Sox3 analysis, embryos were injected at the 8-cell stage into one dorsal animal blastomere with 10 ng of Rspo2 MO or control MO, together with Myr-BFP-HA RNA. For Rspo2 localization studies, embryos were injected at the 32-cell stage into one dorsal animal blastomere with RNAs encoding Rspo2-FLAG, Rspo2-GFP, or Rspo2 deletion constructs, as indicated. For biochemical assays, four-to eight-cell embryos were injected in the animal region with RNAs encoding HA-Vangl2, FGFR1, FGFR4, Rspo2-FLAG, or Rspo2 deletion constructs, as indicated in the figure legends.

### Neural tube defect analysis

Embryos injected unilaterally with Rspo2 MO were cultured until stages 16-17 and fixed for morphological analysis. Neural tube closure phenotypes were scored based on the degree of neural fold convergence and closure on the injected side. Phenotypes were classified as normal, mild, or severe. Mild defects were defined by delayed or incomplete neural fold convergence, whereas severe defects were defined by pronounced failure of neural tube closure. Phenotype frequencies were calculated from the total number of scored embryos per condition. Data shown are representative of two independent experiments.

### Immunostaining and imaging

Embryos were fixed at the stages indicated in the figure legends. For detection of endogenous Vangl2, embryos were fixed in 2% trichloroacetic acid for 30 min and stained with rabbit polyclonal anti-Vangl2 antibody 1:150 as described (Ossipova et al., 2022). Mouse monoclonal anti-Sox3 antibody, 1:100 (Developmental Studies Hybridoma Bank (DSHB, clone DA5H6) was used to assess effect of Rspo2 MO on neural differentiation and HA-tagged lineage tracer (myr-BFP-HA) was detected using rabbit polyclonal anti-HA antibody 1:1000 (Bethyl). For detection of tagged Rspo2 constructs, embryos were fixed in MEMFA solution containing 0.1 M MOPS pH 7.4, 2 mM EGTA, 1 mM MgSO^4^ and 3.7% formaldehyde. GFP-tagged Rspo2 constructs were detected by GFP fluorescence. FLAG-tagged Rspo2 constructs were detected by immunostaining with anti-FLAG antibody (1:300; M2, MilliporeSigma). Phalloidin-A647 (Invitrogen) was used to label filamentous actin and define cell boundaries in Rspo2 localization experiments. Secondary antibodies conjugated to Cy2, Cy3, or equivalent fluorophores were used to detect primary antibodies. Standard negative controls lacking primary antibody were performed to assess background staining and channel cross-reactivity. Neural plates were dissected, put in mounting medium (Vectashield) and imaged *en face* as indicated. Images of whole neural plate explants were acquired using BC43 confocal microscope (Andor), and tiled images were stitched using Fusion software (Andor). Brightness and contrast were adjusted uniformly across comparable images for presentation.

### Quantification of planar polarity

Planar polarity of Vangl2 and Rspo2 was quantified from *en face* neural plate images oriented with the anterior side upward. For Vangl2 polarity analysis, membrane-associated Vangl2 signal was analyzed in control and Rspo2 MO-injected cells. For each cell, a polarity vector was calculated from the cell centroid toward the intensity-weighted centroid of membrane-associated Vangl2 signal. The resulting vector angle was used as the polarity angle for that cell. For Rspo2 polarity analysis, cell outlines were segmented from the phalloidin channel using Cellpose and manually inspected. For each segmented cell, a 2-pixel-wide internal membrane ring was extracted. Rspo2-FLAG intensity was measured within this membrane region. Background intensity was estimated independently for each embryo as the median of the mean membrane intensity of the dimmest 20% of cells. Rspo2-positive cells were defined as cells with mean membrane intensity at least 80% above background. For each Rspo2-positive cell, membrane pixel positions were converted into unit vectors extending from the cell centroid to each membrane pixel. These vectors were weighted by background-subtracted Rspo2 intensity. The weighted vector sum was used to calculate the polarity angle, with 0° corresponding to anterior. Rose plots show raw counts of cells binned by polarity angle. Circular statistics were performed using the Rayleigh test to assess deviation from a uniform angular distribution and the V-test to assess directional bias toward the anterior axis. Custom Python scripts developed with assistance from ChatGPT/OpenAI were used for quantification, plotting, and statistical analysis. All outputs were reviewed by the authors and verified by visual inspection of analyzed cells overlaid on the original images.

### Statistics and data presentation

Phenotype frequencies were calculated from the number of embryos scored in each category. Circular statistics for polarity analyses were performed using the Rayleigh test and V-test. Numbers of analyzed cells and embryos are indicated in the figure legends. Immunoblotting experiments were repeated independently as indicated. Graphs and plots were generated using custom Python scripts, Prism and Microsoft Excel.

### Ectoderm explant preparation

For ectoderm explant experiments, embryos were injected with Rspo2 MO at the four-to eight-cell stage, as indicated. At stage 9, vitelline membranes were removed manually, and animal cap explants were dissected. Explants were cultured in 0.6× MMR until the indicated sibling stage. For analysis of endogenous Vangl2 tyrosine phosphorylation, explants were collected at stage 17 and processed for immunoprecipitation and immunoblotting.

### Immunoprecipitation and immunoblotting

Embryos or ectoderm explants were lysed at the stages indicated in the figure legends in ice-cold lysis buffer supplemented with protease and phosphatase inhibitors as described (Chuykin et al., 2018). Lysates were clarified by centrifugation before immunoprecipitation. For analysis of exogenous Vangl2 phosphorylation, HA-Vangl2 was immunoprecipitated using HA-Trap beads (Chromotek). For analysis of endogenous Vangl2 phosphorylation, lysates from ectoderm explants were immunoprecipitated with rabbit polyclonal anti-Vangl2 antibody (ABN2242, Milliporesigma) and Protein A Sepharose. For co-immunoprecipitation of Rspo2 and Vangl2, Rspo2-FLAG was immunoprecipitated using anti-FLAG agarose (M2, Milliporesigma), and co-immunoprecipitated HA-Vangl2 was detected by immunoblotting. Vangl2 tyrosine phosphorylation was assessed by immunoblotting with anti-phosphotyrosine antibody (pY20, Santa Cruz). Immunoprecipitated Vangl2 was detected with mouse monoclonal anti-HA (12CA5) or anti-Vangl2 (C8, Santa Cruz) antibodies. Expression of injected constructs in lysates was confirmed using anti-HA (12CA5), anti-FLAG (M2, Milliporesigma), anti-GFP (GFP-B2, Santa Cruz). Chemiluminescent signals were detected using a Bio-Rad ChemiDoc imaging system. Representative immunoblots from independent experiments are shown as indicated in the figure legends.

## Acknowledgements

We thank Alice Reis for the initial observation of Rspo2 polarization, and members of the Sokol laboratory for useful discussions. This study was supported by the NIH grant R35 GM122492 to SYS.

